# Sequence-based models for RNA-Protein interactions imputation might be insufficient for novel signal prediction in eCLIP data

**DOI:** 10.64898/2025.12.14.694209

**Authors:** Arsenii Rybakov, Daniil Khlebnikov, Daria Ovchinnikova, Arina Nikolskaya, Arsenii Zinkevich, Andrey Mironov

## Abstract

Predicting specific RNA-protein interactions remains a challenging task: despite the existence of numerous methods, a unified approach has yet to emerge. Additional difficulties emerge from the properties of in vivo IP experiments and their systematic biases, such as the overrepresentation of highly expressed RNAs. Here, we present the PLERIO machine learning framework, which utilizes eCLIP data for a single protein to reconstruct the full spectrum of its potential interactions with the cellular transcriptome (i.e., both highly expressed and lowly expressed RNAs). In an effort to extrapolate our methodology to a multi-protein paradigm for de novo prediction of RNA-protein interactions on proteins lacking available eCLIP data, we extended our approach to 220 cellular proteins. We then demonstrate that this approach might not be well tailored to the limitations of current in vivo immunoprecipitation data and may only be meaningful for in vitro experiments such as RNAcompete.

## 1. Introduction

Interactions between RNA and proteins play a pivotal role in post-transcriptional regulation, alternative splicing, RNA stability, and its intracellular localization [1–3]. It has been observed that certain functions of non-coding RNAs may be facilitated by one or more nuclear proteins or their complexes [4–6]. Notably, this has been documented in for XIST [7], NEAT, MALAT1, and HOTAIR [8]. Conversely, the absence or presence of protein interaction with non-coding RNA (CHD4 [5], PRC2 [6]) has been demonstrated to regulate the functions of certain proteins. Furthermore, proteins have been demonstrated to facilitate RNA interactions with chromatin [9]. The problem of RNA-protein interactions therefore demands careful study to grasp the systems biology of regulatory interactions in the cell.

To date, there are limited data on in vivo RNA-protein contacts. For instance, eCLIP [10] results are available for 120 proteins in the K562 cell line [11], with each dataset dominated by information on highly expressed RNAs. However, RNAs with low expression are of greatest interest for studying regulatory interactions.

Cross-linking immunoprecipitation (CLIP) methods facilitate the mapping of RNA-binding proteins (RBPs) at the transcriptome level in vivo. The immunoprecipitation protocol has long been known to exhibit certain flaws and artifacts [4]. For instance, there is a tendency for highly expressed RNAs to be overrepresented in experimental data compared to controls. This sampling bias is introduced at the protocol level [12,13]. For reasons unrelated to the specificity of their interaction, highly expressed

RNAs clog up more interactions of the assay target protein. Standard data processing protocols, which permit the introduction of control background data into the pipeline, serve to further exacerbate this issue. Even in scenarios where a portion of the experiment’s coverage originates from interactions between lowly expressed RNA and the target protein, there is no way to ascertain that this coverage will surpass the statistical significance threshold necessary for determining the region of this RNA as an interaction peak [14,15] (**Figure 1A**). Additionally, eCLIP data is unable to differentiate between facultative interactions of a protein with RNA regions that are exclusively preferred in cooperative binding due to the single-protein nature of the elementary step of the protocol [16,17].

**Figure 1.**
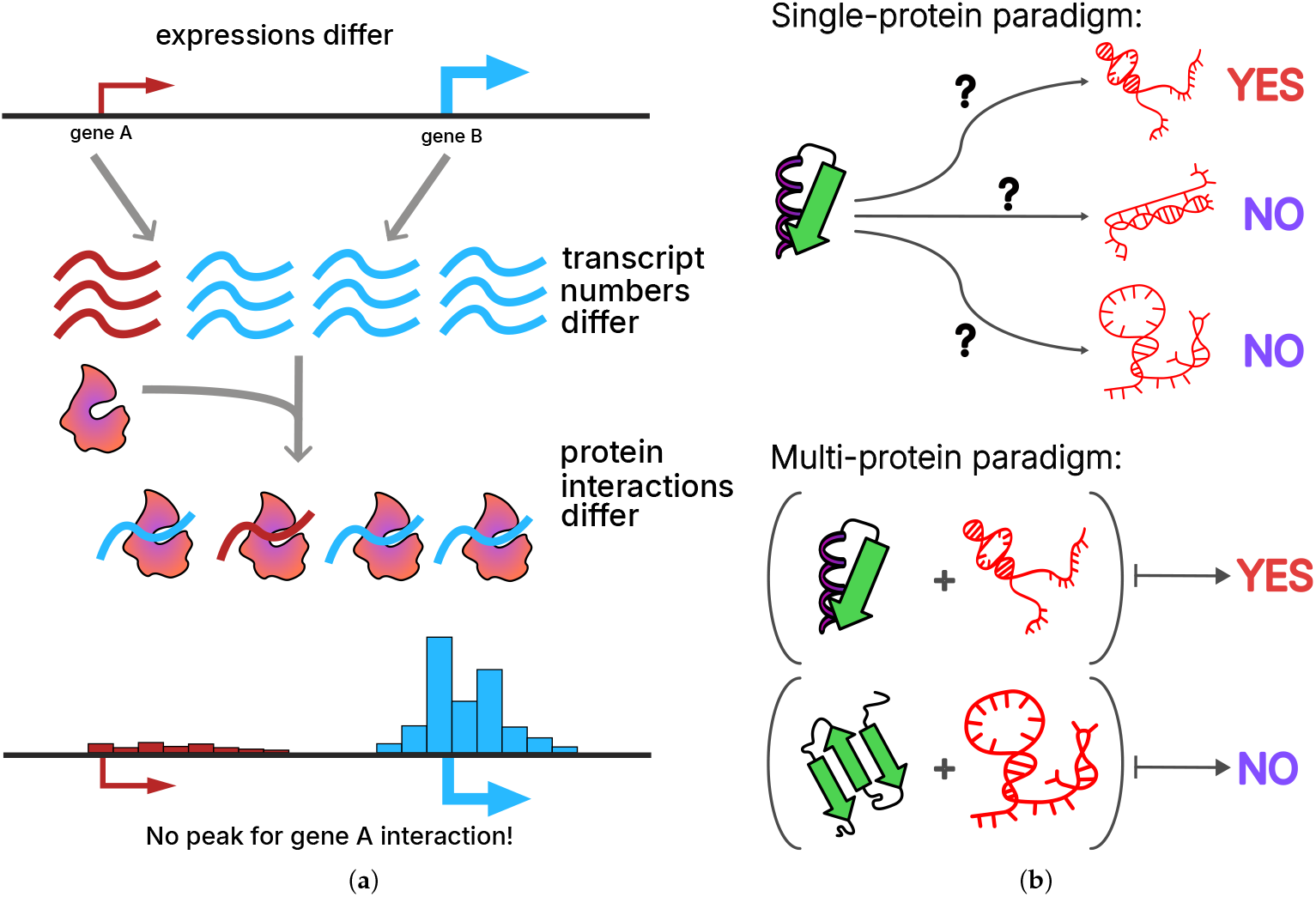
Challenges associated with predicting RNA-protein interactions from in vivo data: **(a)** the systematic bias based on disparities in RNA expression gives rise to the overrepresentation of specific genes in the results; **(b)** the issue of two paradigms: single-protein (top) and multi-protein (bottom) models for predicting interactions.

Furthermore, immunoprecipitation experiment data is also used in the context of predicting RNA-protein interactions de novo [18]. This task is commonly presented in two distinct classes. The first one encompasses machine learning models that have been trained on RNA-protein interaction data for a specific protein. These models predict the probability of RNA-protein contact for a pair of protein and a novel RNA that the model has not previously encountered (see **Figure 1B**, top). The objective of these models is to identify characteristic sequence or structural motifs preferred by a given protein. However, the “one protein - one model” approach is inherently limited in its ability to generalize the patterns of RNA-protein interactions for proteins that were not included in the training data. The utility of these models is primarily limited to supplementing the existing interactome with various semi-extractable or weakly expressed transcripts. Consequently, the utilization of these models in identifying de novo interactions is constrained by the principles of homology.

In order to overcome these limitations, attempts can be made to create multi-protein models that are trained on multiple eCLIP datasets simultaneously (see **Figure 1B**, bottom). In this paradigm, different proteins and multiple RNAs are considered simultaneously, allowing for the use of common binding patterns, knowledge transfer between proteins, and, ideally, imputation of missing interactions. This approach can be applied, for instance, in the context of RNA–protein–DNA triads (or more simply, predicting unknown interactions between a given protein and RNA). For instance, methodologies for employing protein and RNA embeddings (ESM2 [19], UniRep [20]) in the context of joint interactome analysis are available.

A multitude of models have been developed to predict RNA-protein interactions [21–28,28–30]. These models employ diverse approaches within given paradigms. RPI-seq [31] utilizes k-mer counting to generate vector representations of proteins and RNA molecules. The k-mer count vectors of proteins and RNAs are subjected to a series of preprocessing steps and subsequently fed into an SVM or random forest classifier. RPI-seq is trained on a limited dataset derived from PDB RNA-protein complex structures, which limits its capacity for organism-specific predictions. RNACommender [32] is a tool that utilizes a protein sequence and calculates the frequency of specific Pfam domains occurrences. The RNA is vectorized by quantifying subgraphs of the RNA structure predicted by RNAfold [33]. The processing of both vectors is accomplished by linear layers of a neural network with sigmoid activations. The vectors obtained in this manner are subsequently converted into a single value using a trained bilinear form. The sigmoid value of this value is the binding probability.

PrismNet [34] proposes a single-protein approach, where an independent model is constructed for each protein separately. A set of such models enables the enrichment of RNA interactomes based on incomplete experimental data. In summary, RNA sequences are uniformly shortened to equivalent lengths and encoded through the application of one-hot encoding. It is further proposed that an additional channel be utilized to represent the icSHAPE accessibility of each nucleotide. The resulting representation is subsequently fed into a convolutional neural network (CNN) comprising multiple residual blocks and a squeeze-and-excitation block. HDR-Net [35] constitutes an enhanced iteration of PrismNet, characterized by a BERT-inspired representation of nucleotides and a more intricate CNN architecture.

However, despite their numerous advantages, both single-protein and multi-protein approaches are highly dependent on the initial eCLIP data, its quality, specificity, and distribution of statistically significant interaction peaks. Despite the existence of established protocols for joint sequential immuno-precipitation [16], the immunoprecipitation technology itself has been the subject of repeated criticism in recent years [13–15,36,37]. The eCLIP protocol is subject to systematic biases. First, crosslinking is far from uniform across all RNA regions. Second, normalization by transcription level may be incomplete. Third, some of the signals reflect associations of RNA with chromatin-binding proteins or RNA-containing complexes rather than direct binding of the target protein to RNA. In this regard, it becomes more attractive to use data on RNA-protein contacts obtained from in vitro experiments, such as SELEX [38], or its alternative, RNAcompete [39,40], where the RNA chip design is calibrated so that different RNA k-mers are evenly represented across it.

In this study, we explore the limitations of single- and multi-protein models in predicting RNA-protein interactions. In the case of single-protein models, the features extracted from RNA were limited to sequence information obtained from eCLIP data. Furthermore, when developing multi-protein models, we incorporated protein features obtained from ESM2 and UniRep embeddings for our sample. The findings demonstrate the feasibility of predicting novel interactions between proteins and RNAs of genes with low, practically background-level expression by single-protein models. We also demonstrate that the task of constructing multi-protein models for de novo prediction of RNA-protein interactions based on in vivo data may be ill-posed due to the variability of biological aspects of experiments. For instance, the different affinities of the antibodies used to the protein can result in differences in the values of pulldown libraries and, sometimes, control libraries. Furthermore, we demonstrate that the latent space of RNA-protein contact features, which can be determined from in vivo experiments such as eCLIP, does not correspond to that obtained from in vitro experiments (RNAcompete).

## 2. Materials and Methods

We selected the datasets encompassing RNA-protein interactions for 220 proteins for this study. In the interest of comprehensively understanding and generalizing the data, it was essential to select the widest possible range of CLIP experiments. To predict RNA-protein data, the PLERIO (Protein-to-Low-Expressed-RNA Interaction Oracle) framework was developed. The source code for this framework is available in the GitHub repository at github.com/yotterni/PLERIO.

### 2.1. Data and Preprocessing

We obtained the results of the eCLIP experiments conducted by the ENCODE [11] consortium for 120 proteins in the K562 cell line and 100 proteins in the HepG2 cell line (see **Supplementary Table S1** for the identifiers of the ENCODE data used). Because the ENCODE database uses a standardized data processing pipeline, we did not reprocess the data. We took data on statistically significant regions of interaction (IDR-thresholded peaks) between 220 proteins and RNA. Thus, the selected data contains approximately three million unique RNA-protein interaction pairs.

For each ENCODE eCLIP experiment, control datasets (RNA-seq corresponding to the pulldown experiment) in BAM format were selected. RNA expression counts were calculated using featureCounts v.2.0.1. RNAcompete data in the form of 7-mer z-scores were obtained from a recent study [40].

We formalized the task of predicting RNA-protein interactions as a binary classification problem. Therefore, the ENCODE data required additional preprocessing. The eCLIP experiment data only contain positive examples of interacting RNA-protein pairs. To train the model, we generated negative examples (non-interacting pairs) as follows: for each protein with known eCLIP results, negative examples were constructed as pairs of the corresponding protein and RNA regions that were not identified as interacting with that protein during the experiment. Thus, either intergenic regions, gene deserts, and introns or regions of RNA that interacted with the protein but for which no peak was detected were selected for the negative sample. To achieve a 1:1 ratio of positive to negative examples in the training sample, a portion of the negative class instances was randomly selected.

The RNA components of RNA-protein pairs were reduced to a uniform size of 200 nucleotides. This fixed input size is required for machine learning models and architectures and helps to avoid overfitting to RNA length. In order to augment the size of the training sample, a regularization technique was employed in tandem with model augmentation for both positive and negative RNA sequences. A 200-nucleotide window shift to all possible positions from -15 bp downstream to +15 bp upstream was incorporated for positive class, and to all possible positions from -15 bp downstream to +15 bp upstream in increments of 5 bp for negative class.

Given that preprocessed RNA data is a fixed-length sequence, it is logical to employ standard natural language processing techniques for RNA vectorization, such as recurrent neural networks or transformers. However, it should be noted that such models are often trained on sizable datasets. In contrast, the we utilize approximately 12,000 examples (including augmentations) for each protein, a substantially smaller dataset. Consequently, a more elementary approach was employed, entailing the quantification of the frequencies of all k-mers in RNA (**Figure 2A**). This approach is reasonably interpretable from a biological perspective [41]. It is well-established that numerous proteins bind to specific RNAs by recognizing short motifs within them. Thus, our proposed vector representation can be regarded as the frequencies of all possible similar patterns of a certain length.

**Figure 2.**
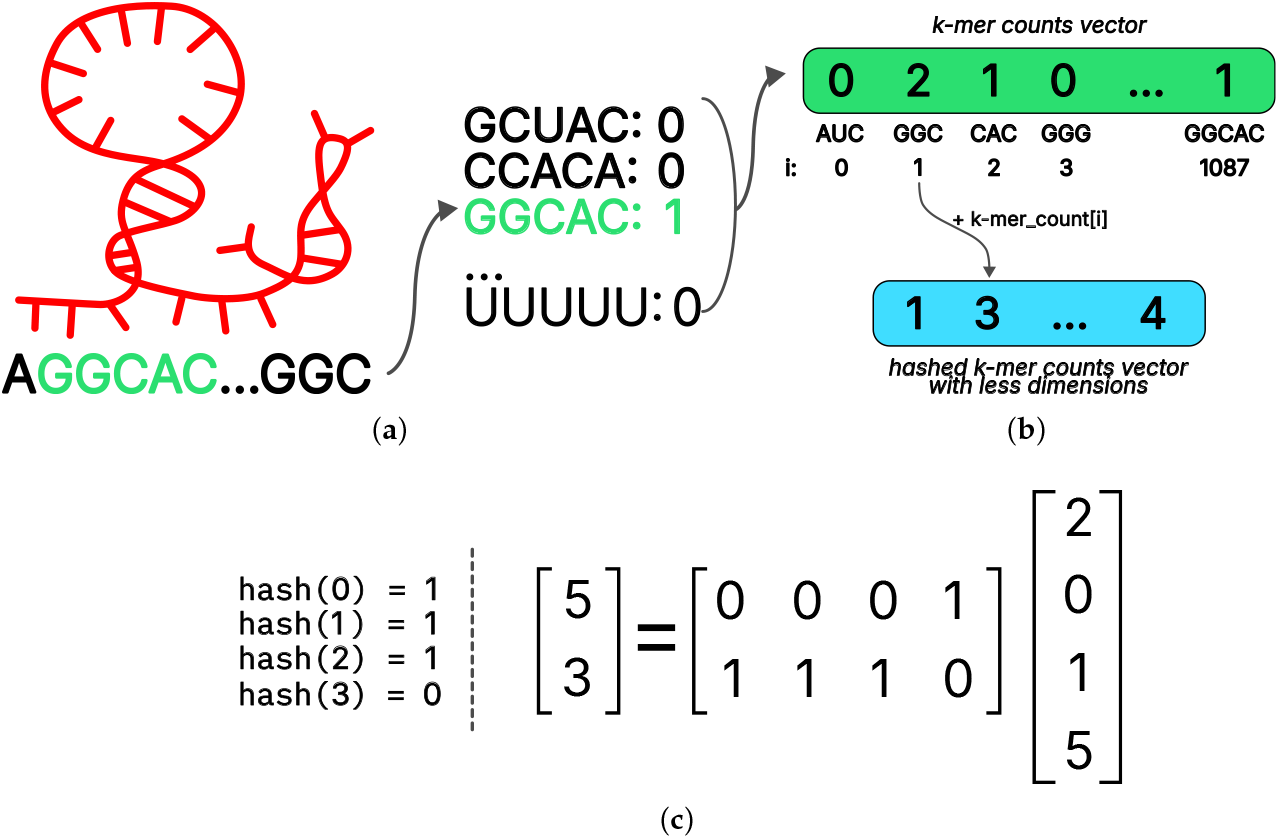
Preprocessing approaches used in the PLERIO framework: **(a)** calculation of RNA k-mer counts, **(b)** vector hashing to reduce the dimensionality of the input data, **(c)** an example of reduction operator linearization for the dimensionality of the input data 4 and the output data 2.

The k values for k-mers length were selected to be 3 and 5. The frequency vectors of 3-mers and 5-mers are concatenated, resulting in a vector representation of RNA with a dimension of 4^3^ + 4^5^ = 1088.

The selection of sequence size is substantiated by the observation that the models exhibited optimal performance on the validation sample within the specified parameters, while the embedding size remained reasonable.

While the calculation of k-mer frequencies as a means of representing RNA has been demonstrated to be a robust regularization for the model, the dimensionality of the resulting embedding is found to be excessively large. When fed into a linear layer with an output dimension that is half the size of the input, the number of model parameters will exceed the size of the training sample. This will cause the model to overfit. To mitigate this issue, we employed a hashing method commonly utilized in natural language processing [42] to reduce the dimensionality of the k-dimensional vector.

The first step in the hashing procedure is determining the desired final dimensionality to be obtained at the conclusion of the procedure. This is followed by the selection of a suitable hash function. In this study, a function from the linear universal family was chosen. Subsequently, for the *i*-th coordinate of the input vector, the value of the hash function *hash*(*i*) was calculated, and the value of the output vector was incremented in this index, thus, *out*(*hash*(*i*))+ = *kmer*(*i*) (**Figure 2B**).

For a fixed hash function, all its values on numbers from 0 to 1087 (as the number of all 3- and 5-dimensions used in the model) can be precalculated in advance. Furthermore, it is feasible to express the hashing procedure in closed form by employing a column-wise stochastic linear operator. An illustration of this procedure, with input and output dimensions of 4 and 2, respectively, is presented in **Figure 2C**. Stochasticity in this context is a property of the operator matrix, indicating the sum of the values in each column is equal to one. Representing hashing as a simple linear transformation facilitates efficient implementation and enables differentiation by the input vector, a property that can be advantageous for interpreting models built on top of hashing. The ESM2 model [19] was selected as the vector representation of the protein due to the fact that models based on Unified representation [20] embedding demonstrated lower quality.

### 2.2. Machine learning models

The single-protein paradigm requires training models for each protein. To accomplish this, we selected and preprocessed eCLIP data from the ENCODE database. A model for a single protein receives the features obtained during RNA preprocessing as input and outputs the probability of interaction between that RNA and the corresponding protein (**Figure 3A**). Such models are necessary for imputing interactions involving lowly expressed RNAs, because it has been shown that information about such RNAs is underrepresented in eCLIP data. eCLIP results are dominated by highly expressed RNAs (see **Figure 4A** in **Results** section), and the statistical processing of experimental data does not include normalization to the level of RNA expression.

**Figure 3.**
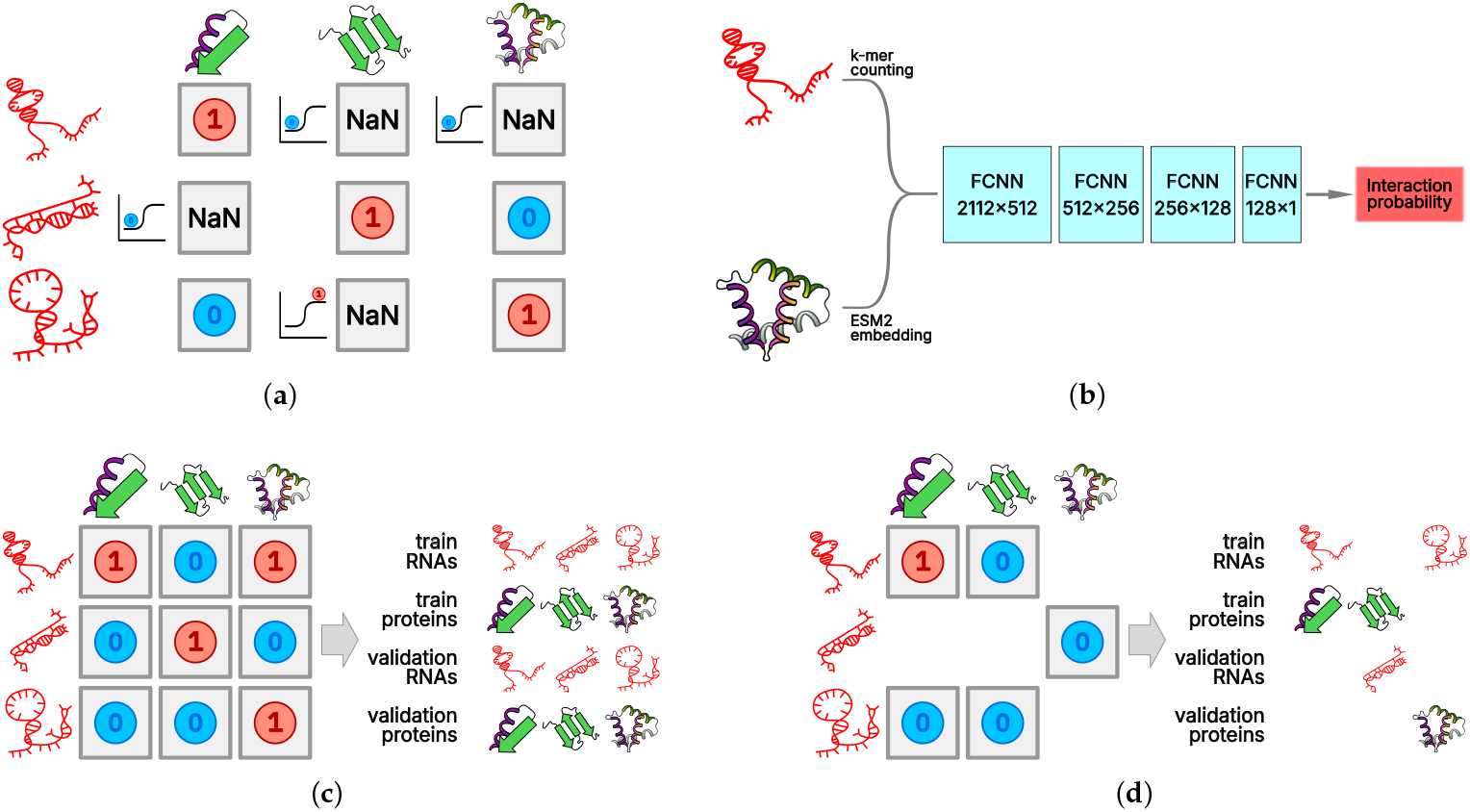
PLERIO framework model data handling. **(a)** single-protein paradigm train-test split, **(b)** multi-protein paradigm model archivtecture, **(c, d)** multi-protein paradigm: naive train-test split **(c)** and the one implemented in this study **(d)**.

**Figure 4.**
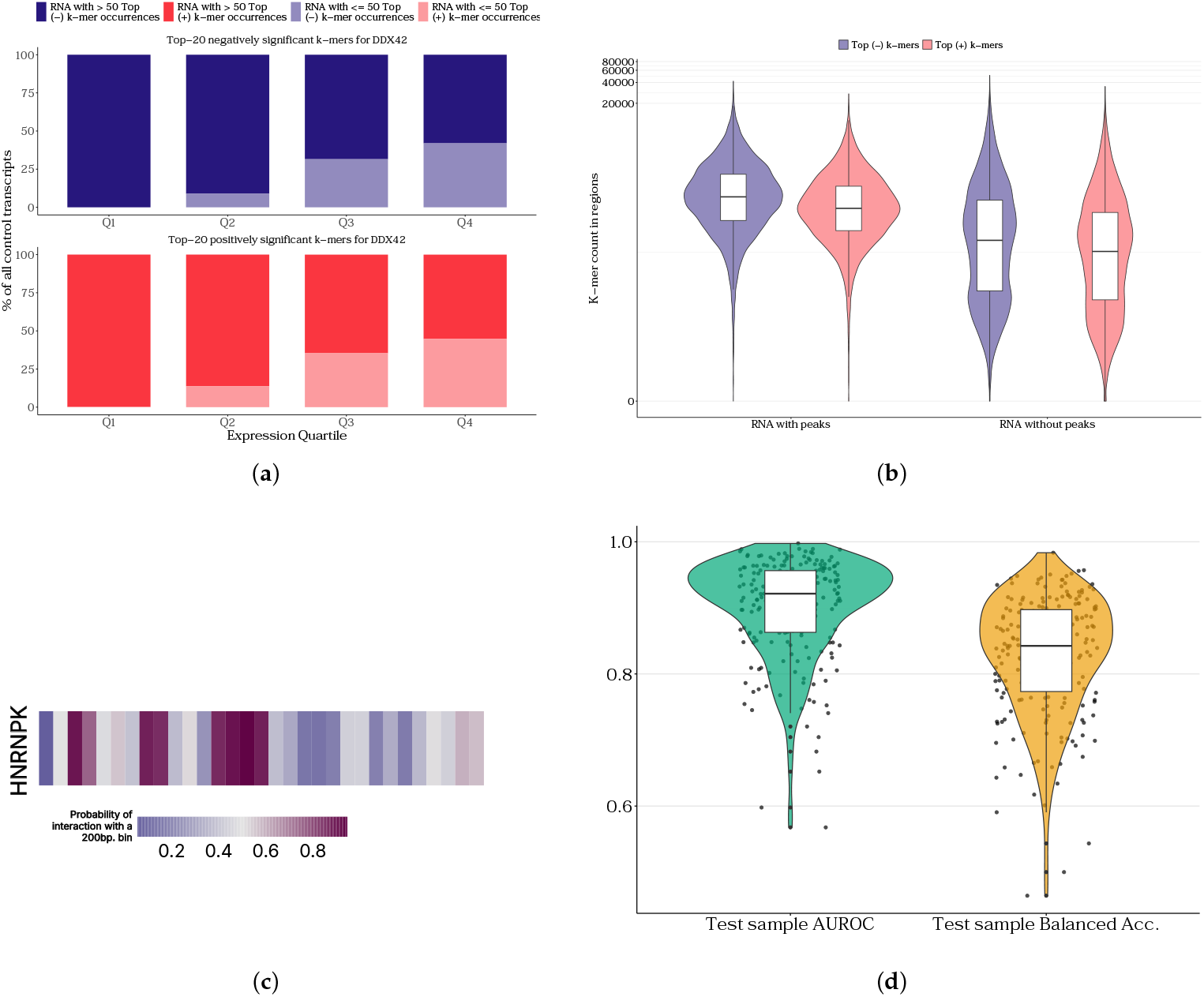
Results of single-protein models developed in the PLERIO framework: **(a)** enrichment of significant k-mers identified by the model in highly and lowly expressed transcripts, **(b)** enrichment of k-mers significant to the model in transcripts determined inside and outside of peaks of interaction with the protein, **(c)** An example of a predicted interaction track for lowly expressed RNA LINC01144 and HNRNPK protein **(d)** quality metrics (AUROC and balanced accuracy) distribution for single-protein models.

For each protein, the set of unique RNAs was randomly divided into training, validation, and test samples. This division was performed independently for the set of unique proteins and RNAs. For a multi-protein model, the protein set was randomly divided. To split the RNA set, first the set of all chromosomes was split, and then the RNAs were divided according to their chromosomes of origin. This approach prevents data leakage from the validation and test samples into the training sample, which can occur when the set of unique RNAs is randomly divided into three samples. Since only a fixed-size vector representation of RNA is fed into the single-protein model, ordinary logistic regression is sufficient.

Creating a multi-protein model involves training a single model using RNA and protein features to predict the probability of interaction between an arbitrary RNA and protein. This approach is necessary primarily for predicting the RNA interactome of proteins for which no experimental data are available. The division of the set of RNA-protein pairs appears to be more complex. With a naive random division of unique pairs into three groups, significant data leakage occurs from the training sample into the validation and test samples. To more rigorously evaluate the model, it must receive not just new RNA-protein pairs, but pairs of RNAs and proteins that did not occurin the training and test samples separately. Otherwise, data leakage occurs, resulting in an overestimation of the model’s quality (**Figure 3C**).

The solution to the aforementioned problem is to split the sets of unique RNAs and proteins independently into training, validation, and test samples. For example, to construct the training sample, RNA-protein pairs should be drawn from pairs of “training” RNAs and proteins. This approach results in partial data loss (**Figure 3D**). Here, we divided the RNAs into three samples using samples of the chromosomes of their origin and randomly divided the set of unique proteins into three groups. Since protein language models accept fixed-size vector representations as input, the vector representation of RNA can be concatenated with it and fed into a fully connected neural network (**Figure 3B**).

### 2.3. Other comparative analyses

In order to make a comparison between in vitro and in vivo data on RNA-protein interactions, a strategy was adapted from a recent study that predicted orthologous RNA-protein interactions in the eukaryotic tree of life [40]. In summary, we reproduced the RNAcompete data processing workflow, constructing a joint embedding matrix of proteins and RNAs followed by SVD with the objective of explaining the RNA profiles matrix norm part of the joint protein-RNA matrix. Furthermore, a comparative analysis was conducted between the latent spaces derived from in vitro and in vivo experimental data using a similar procedure for eCLIP data. The z-score profile of 7-mers from RNAcompete was substituted with a metric derived from peaks. For each 7-mer, we derived its binding signal by taking the maximum value of fold change times − log10(*p* − *value*) across all eCLIP peaks where that 7-mer was present. This maximum-value approach was chosen to avoid signal averaging effects that could mask genuine binding preferences, as averaging would dilute specific interactions with numerous weaker, non-specific binding events commonly observed in protein-DNA interactions. Subsequently, the 7-mer vector was centered and normalized within a single protein, thereby yielding a proxy z-score. Protein embeddings for such a matrix were obtained using methods similar to those in the original study: HMMscan [43] was used to find and then extract target RNA-binding domains from the proteins, created RNA- and protein-embedding matrices were decomposited. See extended list of RNA-binding domains used for this study in **Supplementary Table S2**.

These and other comparative analyses were conducted using in-house Python and shell scripts. Visualization was performed in R. To reproduce the analysis, one can find the scripts in the accompanying GitHub repository at github.com/dkhlebn/PLERIO_scripts.

## 3. Results

The models and their weights obtained during the development of the PLERIO framework are available on the GitHub repository for 220 studied proteins.

### 3.1. Single-protein models performance

The primary objective of generating single-protein models is to enhance the RNA interactomes of proteins with existing eCLIP data for lowly expressed RNAs. For each protein, the resulting model was interpreted in the context of k-mers that are significant for predicting the existence or absence of potential interactions with RNA. A comparison was made of the frequency of k-mers significant for the model in highly expressed and low-expression transcripts (**Figure 4A**, with DDX42 protein model shown as a chosen example based on top model performance) and of the presence of such k-mers in statistically significant peaks of protein-RNA interactions (**Figure 4B**, also see **Supplementary Figure S1** for stratified regions and gene length-normalized analysis). It has been demonstrated that RNA molecules with lower expression levels frequently contain a greater number of k-mers that are critical for model prediction. However, the presence of these RNAs in peaks is not guaranteed, as their coverage in the immunoprecipitation experiment is subject to systematic bias due to the lack of normalization for expression levels. Despite the incorporation of background expression level normalization in the ENCODE consortium’s processing pipeline, our findings indicate that this may be insufficient. This effect, though less pronounced, is also conserved when applying normalization by RNA length (see **Supplementary Figure S2**).

We provide examples of model predictions for some proteins on specific lowly expressed RNAs (see **Figure 4C**; for other examples of different proteins and RNAs, see **Supplementary Figure S3**). For these, we hand-picked lowly expressed RNAs from a sample of proteins RNA-seq experiment data and ran the inference for them. Given the potential variability in RNA length and the capacity of the model to only handle sequences up to 200 nucleotides, a screening approach is employed. The model systematically traverses each RNA, with a 200-nucleotide window and a 50-nucleotide step. This approach is more reliable than dividing the RNA sequence into independent segments of 200 nucleotides, as it allows for the spatial proximity of neighboring windows to be taken into account. We note that some of the examples we present fall into peaks of select proteins (e.g., HNRNPK-LINC01144 RNA interaction). This is not the case for all of the examples we present.

The 220 models constructed (120 for the K562 cell line and 100 for HepG2) yielded a distribution of prediction quality metrics for RNA-protein interactions in the single-protein model training paradigm (**Figure 4D**). The quality of our model, which bases its predictions solely on RNA sequence data, surpasses existing methods for predicting RNA-protein interactions based on deep learning models. This is supported by the observation that the median AUROC distribution of our models exceeds that of competing approaches (**Table 1**). That said, our proposed approach is also much more straightforward to implement and interpret. The models we have compared the quality of predictions against utilize more complex approaches to working with RNA sequences, namely convolutional neural networks (PrismNet) and a composition of a convolutional network with a transformer (HDRNet). Furthermore, it appears that the positive-to-negative class balance in the training sample is also a significant factor. Our models were trained on samples where the ratio of positive and negative class samples was 1:1, while PrismNet’s appears to be close to 2:1.

**Table 1.**
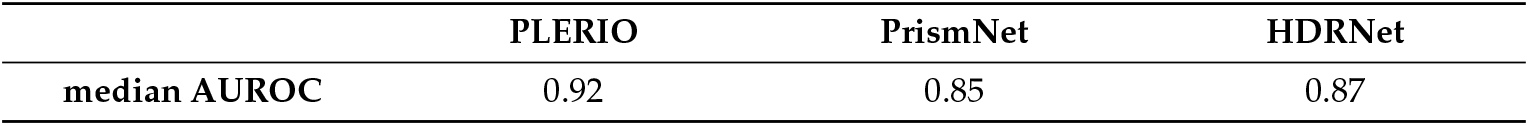
Median values of the AUROC metric for single-protein classifiers of RNA-protein interactions of PLERIO framework compared against previous approaches. PrismNet and HDRnet metrics were retrieved from the models source publications.

The k-mer hashing approach we employed enables us to reduce the embedding dimension of RNA k-mer frequencies by up to tenfold without sacrificing quality. Furthermore, using different output embedding sizes did not significantly improve the quality of the predictions for any of the tested proteins. (see **Supplementary Figure S4**). The relative success of hashing raises the question of whether we can effectively compress even larger vectors. Thus, we plan to use hashes to compress k-mer frequency vectors with *k* > 5 in the future.

### 3.2. Multi-protein model performance

Another objective of the study was to evaluate the feasibility of developing a unified model to predict RNA-protein contacts de novo for proteins for which in vivo immunoprecipitation data has not been produced or published due to various reasons. Creating such a model is of great value, both in itself and for applied tasks. One example is integrating model predictions with various types of pairwise interaction data [9], such as DNA-protein ChIP-seq data or RNA-chromatin interactome data (Red-C [44], RADICL-seq [45], and GRID-seq [46–48]).

To evaluate the specificity of the multiprotein model, we created binding probability tracks for 18 proteins that have various cellular functions and for which eCLIP data is unavailable. **Figure 5A** shows interaction tracks for selected proteins without publicly available eCLIP data and MALAT1 RNA (see **Supplementary Figure S5** for interaction tracks with several different RNAs). Unfortunately, the model’s inability to generalize parameters for de novo imputation of RNA-protein interactions is evident. Perhaps the areas of non-specific activation of the model on MALAT1 for many proteins are regions that could potentially interact electrostatically with proteins, e.g., GC tracts [17]. As expected for a randomly selected set of proteins and random non-coding RNA, no specific binding sites on MALAT1 are observed for all 18 proteins. With corresponding metric values of balanced accuracy (0.79), AUROC (0.87), and MCC (0.52), the model designed in a multi-protein paradigm appears to generate a large number of false positives. This behavior is most likely caused by a scarcity of training data. The model was trained using K562 data with only 90 proteins in the training sample, and 30 were set aside for validation and testing.

**Figure 5.**
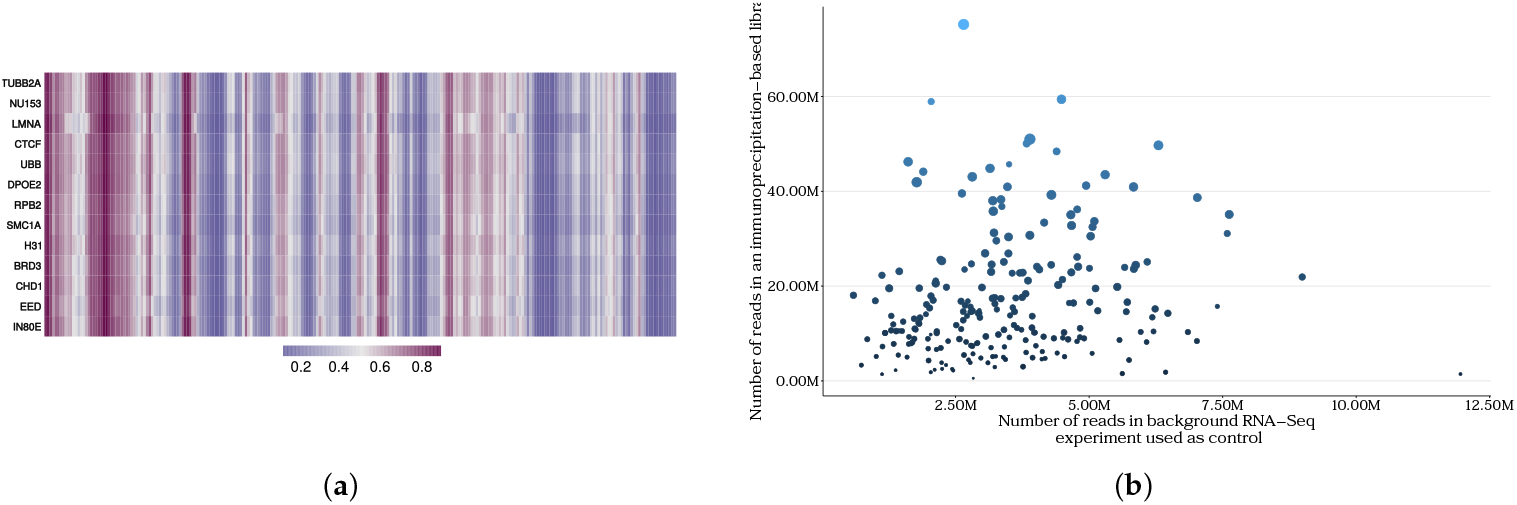
The PLERIO framework multi-protein model: **(a)** an example of tracks of several proteins on MALAT1 RNA; **(b)** the difference in library sequencing depth and the number of statistically significant RNA-protein interaction peaks in eCLIP data for the proteins in the training sample, size and color of a point correspond to the eCLIP library size.

Moreover, a critical component in the construction of this model involves the consideration of the equivalence of the data utilized for its training. The inherent heterogeneity of the biophysical characteristics of eukaryotic proteins presents a challenge when conducting immunoprecipitation of their complexes with RNA, as antibodies for those proteins may exhibit varying affinities. This phenomenon results in disparities in the depth of pulldown sequencing libraries among diverse proteins, consequently yielding varying amounts of training data for different proteins (see **Figure 5B**). Furthermore, recent studies have raised questions about the biological purity of immunoprecipitation data, and attempts have been made to correct for biological artifacts in such experiments [13,36,37]. As a result, the issue of predicting de novo RNA-protein interactions based on available in vivo eCLIP data alone is either ill-posed or remains infeasible due to the limited amount of data currently available.

### 3.3. eCLIP and RNAcompete consistency

One emerging domain of research within the broader framework of RNA-protein interactome studies involves the study of in vitro methods and extrapolating their findings to the context of in vivo scenarios. One of the most prevalent in vitro methods is RNAcompete. A thorough analysis of the experimental data has already demonstrated its effectiveness in determining the specificity of binding to motifs in RNA sequences for proteins with or without an RNA-binding domain [17], and has also contributed to the expansion of the range of RNA-protein interactome motifs across the eukaryotic tree of life [40]. In order to determine the viability of obtaining such results through in vivo experimentation, a similar analysis was conducted on a joint embedding matrix of proteins and their interactomes that was obtained from eCLIP data.

In JPLE, the RNA interactome of each specific protein was represented as a vector of 7-mer z-scores retrieved from RNAcompete. We selected a subset of the proteins from the original study for which a public set of eCLIP peaks also existed. As means to replace the RNAcompete z-score vector, the average signal was normalized across the entire set of 7-mers for those peaks in which each specific 7-mer was present. We then reproduced the singular value decomposition (SVD) analysis of both combined matrices (RNAcompete and eCLIP data) for these proteins.

We demonstrate that RNAcompete data provides a more rapid explanation of the variance in RNA-protein interaction data (**Figure 6A**). The decomposition curve of the RNAcompete-based matrix reaches a plateau faster than than that of the eCLIP data. Furthermore, we investigated the latent embeddings spaces into which SVD projects the corresponding matrices. It was observed that the distributions of cosine distances between proteins in these spaces differ between the one constructed based on RNAcompete and the one obtained from eCLIP (**Figure 6B-C**, wasserstein’s W1 distance 0.1029).

**Figure 6.**
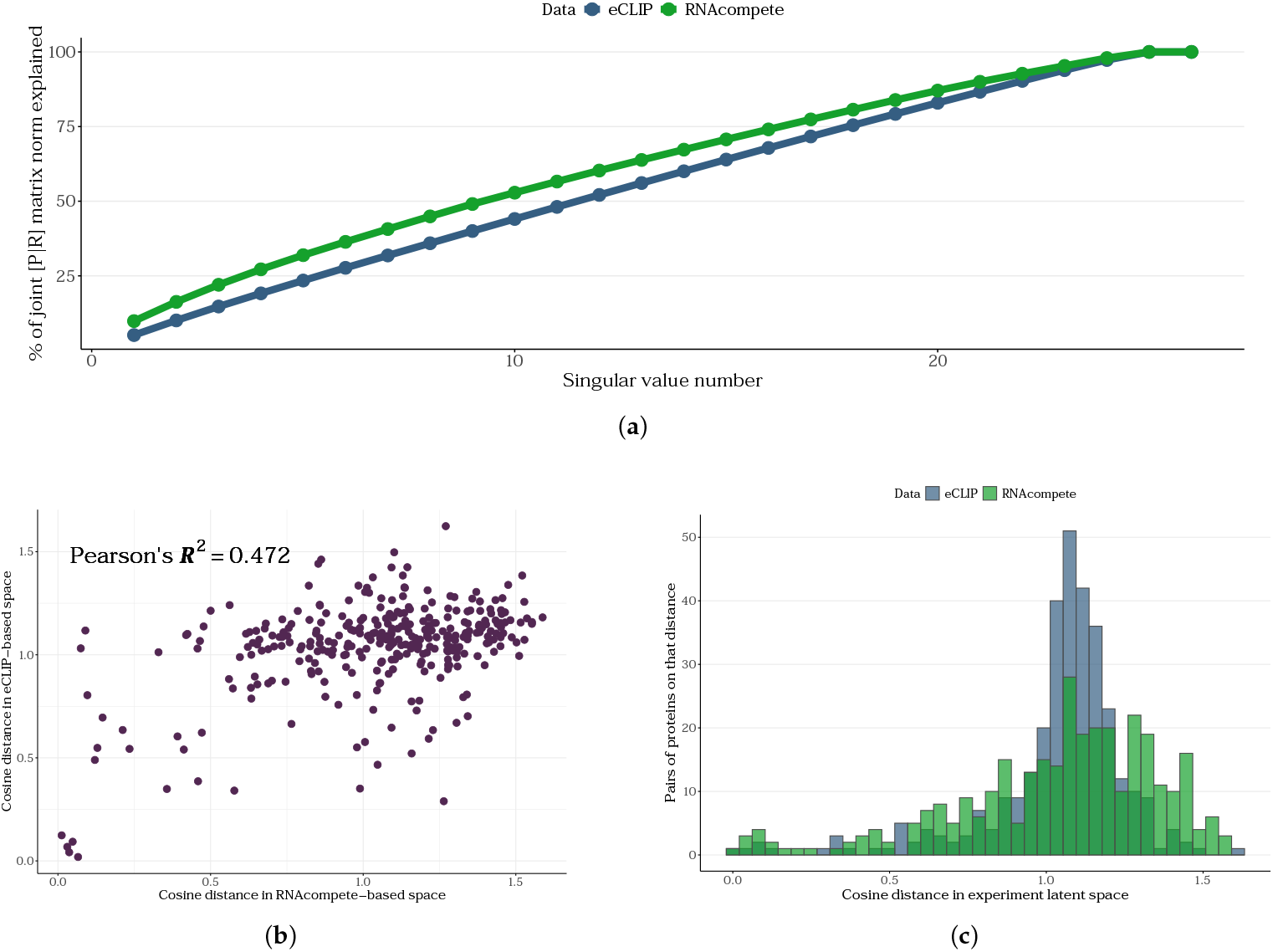
A comparison of the JPLE approach for RNAcompete and eCLIP data is shown below: **(a)** matrix norm proportion explained by the JPLE algorithm for proteins with both eCLIP and RNAcompete data, **(b)** distribution of distances in two spaces, based on RNAcompete and eCLIP, pearson’s *R*^2^ = 0.472, *p*-value < 10^−16^, **(c)** same, within each of the spaces.

The sample of proteins in the original JPLE study relied on selecting proteins with canonical RNA-binding domains. Upon incorporating proteins with RNA-binding domains that were not previously considered in JPLE into the specified space, it was observed that the distance distribution remained unchanged (**Supplementary Figure S6**). Thus, we demonstrate that the underlying biology of in vitro and in vivo experiments differs and reflects the faults of eCLIP-based data. Given the mounting criticism in the literature regarding the eCLIP protocol, as well as the results reported in this study, it is essential to conduct a thorough analysis to ensure the applicability of eCLIP.

## 4. Discussion

The identification of functional RNA-protein interactions constitutes a pivotal aspect of the investigation of non-coding RNA biology and eukaryotic gene regulatory networks. The expansion and refinement of RNA interactomes of nuclear proteins for which public datasets of in vivo experiments exist is of particular interest. It is evident that protocols designed to identify statistically significant interactions between target proteins are marred by a multitude of experimental (technical) and biological flaws, a fact that has already been recognized and documented within the community. In this study, we critically evaluate the potential for expanding interactomes through the incorporation of low-expression RNAs, which exhibit inadequate coverage in immunoprecipitation pulldown experiments. Additionally, we explore the development of a comprehensive RNA-protein binding model that facilitates de novo prediction of such interactions. This model is particularly essential for the imputation of data concerning proteins that are not represented in public databases due to a lack of experimental data.

In this study, PLERIO, a newly developed framework, was utilized to illustrate the successful incorporation of low-expression RNAs into the RNA-protein interaction interactome within the single-protein model paradigm. The proposed model, which exclusively utilizes RNA sequence properties, has demonstrated superior performance in comparison to previously developed approaches that have employed more intricate neural network architectures. We demonstrated that the adoption of particular preprocessing methodologies, such as k-mer hashing [42], can be utilized to simplify the training process of the models.

The interchangeability of promising results obtained from in vitro methods and in vivo immuno-precipitation data was also examined. The results demonstrate that the sequence data obtained from eCLIP defines a protein embedding space that differs from that defined by RNAcompete data. This way, we demonstrate that RNA and protein sequence data alone may be insufficient for single base-resolution de novo prediction of RNA-protein interactions from in vivo data. When constructing the model, we deliberately excluded features related to RNA and protein structure. Moreover, we did not account for the cooperativity of protein binding to different transcripts when constrained by other nuclear proteins. We propose that sequence properties may not be the key to predicting interactions de novo because, in contrast to the symmetric task of predicting TF interactions with DNA, the mutual arrangement of RNA and protein pockets in space plays a much more crucial role. Additionally, the function of intrinsically disordered regions (IDRs) within proteins in the context of RNA-protein interactions should be studied. It has been established that a considerable number of nuclear regulatory proteins are endowed with IDRs or are characterized by a complete absence of structure [49,50]. This, in turn, creates another degree of freedom for RNA bound to IDR-containing proteins. Current immunoprecipitation protocols do not guarantee that the regions of RNA where protein binding occurs are true binding sites [15], because interaction peaks in such experiments are often determined by GC-rich sequences due to their electrostatic properties [17]. Future efforts should integrate structural, spatial, and biophysical information, alongside high-resolution experimental datasets, to achieve a more complete and mechanistic understanding of RNA-protein interactomes. Such integrative approaches will be essential for accurately modeling the complexity of nuclear regulatory networks and for extending predictive frameworks to proteins that remain underrepresented.

## Supplementary Materials

The following supporing information is included: **Supplementary Information**.

## Author Contributions

Conceptualization, A.M. and D.Kh.; methodology, A.R., D.Kh., A.Z. and A.N.; software, A.R., D.Kh., D.O.; formal analysis, A.R., D.Kh, D.O.; writing—original draft preparation, D.Kh.; writing—review and editing, D.Kh.; visualization, A.R., D.Kh. and D.O.; supervision, A.Z, A.N. and A.M.; project administration, A.Z., A.M.; funding acquisition, A.M. All authors have read and agreed to the published version of the manuscript.

## Data Availability Statement

The data that support the findings of this study are available from the corresponding author, D.Kh., upon reasonable request.

## Acknowledgments

The authors thank Anastasia A. Zharikova for their valuable comments on manuscript.

## Disclaimer/Publisher’s Note

The statements, opinions and data contained in all publications are solely those of the individual author(s) and contributor(s) and not of MDPI and/or the editor(s). MDPI and/or the editor(s) disclaim responsibility for any injury to people or property resulting from any ideas, methods, instructions or products referred to in the content.

