## Supplementary Information for "Sequence-based models for RNA-Protein interactions imputation might be insufficient for novel signal prediction in eCLIP data"

### Supplementary Materials


| | | |
|---|---|---|
| Table S1: | ENCODE dataset identifiers for data used in the study. | 2 |
| Table S2: | Extended RNA-binding domains list used for this study. | 9 |
| Figure S1: | Stratified analysis of DDX42 model k-mer importance correlation with the k-mer being the statistically significant peak of the DDX42-RNA interaction: **A** figure from original manuscript; **B** same, but the number of important k-mers was normalized to total region length on a transcript; **C** number of important k-mers on transcripts with or without peaks on them, if and RNA has a peak on it, it is split into peaks and regions outside the peaks; **D** same, but normalized to the according region's legth transcript-wise. | 11 |
| Figure S2: | Enrichment of significant k-mers identified by the model based for DDX42 protein in highly and lowly expressed transcripts, normalized by gene length. | 11 |
| Figure S3: | Examples of single-protein models ncRNA-protein contact inference in PLERIO framework: **A** CPEB-DT and CPEB4 protein, possible family-wise regulation; **B** THOC7-AS1 and NCBP2 protein, possibly connected via RNA export; **C** GLIS2-AS1 and RBFOX2 protein, possibly connected through involvement in splicing control; **D** NDUFB2-AS1 RNA and SAFB and SAFB2 proteins, possible antisense RNA interaction; **E** PRKAR2A-AS1 and NONO protein, possibly interacting in paraspeckles; **F** RN7SL23P RNA and SSB and TROVE2 proteins. | 12 |
| Figure S4: | The effects of varying k-mer hasing output vector dimension on model performance. | 13 |
| Figure S5: | Examples of multi-protein models ncRNA-protein contact inference in PLERIO framework: **A** NEAT1; **B** HOTAIR; **C** TERC; with multiple proteins covering a wide range of functions where RNA-binding might not be needed. Yet, the interaction profile is practically the same. | 14 |
| Figure S6: | Distribution of cosine distances in eCLIP-based latent space obtained by JPLE algorithm on available proteins with a canonical RBD or a domain that can be less strictly considered RNA-binding. Wasserstein's W1 distance between two distributions is 0.01697. | 15 |

**Supplementary Table S1.** ENCODE dataset identifiers for data used in the study.

| Protein | Cell line | eCLIP ENCODE accession |
|---|---|---|
| AARS | K562 | ENCFF128BDG |
| AATF | K562 | ENCFF955LYD |
| ABCF1 | K562 | ENCFF941GJK |
| AGGF1 | HepG2 | ENCFF992LCW |
| AGGF1 | K562 | ENCFF810NKV |
| AKAP1 | HepG2 | ENCFF749IUB |
| AKAP1 | K562 | ENCFF033EFV |
| AKAP8L | K562 | ENCFF205UTG |
| APOBEC3C | K562 | ENCFF472CCL |
| AQR | HepG2 | ENCFF323PUC |
| AQR | K562 | ENCFF949QGM |
| BCCIP | HepG2 | ENCFF134LIU |
| BCLAF1 | HepG2 | ENCFF810EUA |
| BUD13 | HepG2 | ENCFF011NAP |
| BUD13 | K562 | ENCFF262ILV |
| CDC40 | HepG2 | ENCFF027MEO |
| CPEB4 | K562 | ENCFF665BGJ |
| CPSF6 | K562 | ENCFF635NZZ |
| CSTF2 | HepG2 | ENCFF349SVY |
| CSTF2T | HepG2 | ENCFF141DOI |
| CSTF2T | K562 | ENCFF878VVA |
| DDX21 | K562 | ENCFF378OYA |
| DDX24 | K562 | ENCFF537KWB |
| DDX3X | HepG2 | ENCFF777NMK |
| DDX3X | K562 | ENCFF087MAJ |
| DDX42 | K562 | ENCFF399GGH |
| DDX51 | K562 | ENCFF799ERK |
| DDX52 | HepG2 | ENCFF005GER |
| DDX52 | K562 | ENCFF198ULC |

See next page

| Protein | Cell line | eCLIP ENCODE accession |
|---|---|---|
| DDX55 | HepG2 | ENCFF797QRS |
| DDX55 | K562 | ENCFF462WAI |
| DDX59 | HepG2 | ENCFF899AWZ |
| DDX6 | HepG2 | ENCFF614NUP |
| DDX6 | K562 | ENCFF904KTV |
| DGCR8 | HepG2 | ENCFF193URO |
| DGCR8 | K562 | ENCFF023YVD |
| DHX30 | HepG2 | ENCFF663QIZ |
| DHX30 | K562 | ENCFF128AKC |
| DKC1 | HepG2 | ENCFF633LQC |
| DROSHA | HepG2 | ENCFF854BHP |
| DROSHA | K562 | ENCFF797BHU |
| EFTUD2 | HepG2 | ENCFF796RMF |
| EFTUD2 | K562 | ENCFF862ZPV |
| EIF3D | HepG2 | ENCFF056UAW |
| EIF3G | K562 | ENCFF634UAC |
| EIF3H | HepG2 | ENCFF813JIK |
| EIF4G2 | K562 | ENCFF207BXK |
| EWSR1 | K562 | ENCFF607ZRF |
| EXOSC5 | HepG2 | ENCFF862EKD |
| EXOSC5 | K562 | ENCFF858CSS |
| FAM120A | HepG2 | ENCFF208ZMI |
| FAM120A | K562 | ENCFF158AOV |
| FASTKD2 | HepG2 | ENCFF498IGK |
| FASTKD2 | K562 | ENCFF423SOR |
| FKBP4 | HepG2 | ENCFF957AOZ |
| FMR1 | K562 | ENCFF480NUZ |
| FTO | HepG2 | ENCFF351ZDU |
| FTO | K562 | ENCFF924OVC |

See next page

**Supplementary Table S1** – continued from previous page

| Protein | Cell line | eCLIP ENCODE accession |
|---|---|---|
| FUBP3 | HepG2 | ENCFF466UMQ |
| FUS | HepG2 | ENCFF972DFZ |
| FUS | K562 | ENCFF861KMV |
| FXR1 | K562 | ENCFF366UJP |
| FXR2 | HepG2 | ENCFF112MVG |
| FXR2 | K562 | ENCFF315HYU |
| G3BP1 | HepG2 | ENCFF899HHP |
| GEMIN5 | K562 | ENCFF050VVI |
| GNL3 | K562 | ENCFF048RLZ |
| GPKOW | K562 | ENCFF674BUM |
| GRSF1 | HepG2 | ENCFF929AWR |
| GRWD1 | HepG2 | ENCFF159MMZ |
| GRWD1 | K562 | ENCFF327DLL |
| GTF2F1 | HepG2 | ENCFF002HYO |
| GTF2F1 | K562 | ENCFF278ZCU |
| HLTF | HepG2 | ENCFF048FDF |
| HLTF | K562 | ENCFF406MSI |
| HNRNPA1 | HepG2 | ENCFF797GSK |
| HNRNPA1 | K562 | ENCFF392AEV |
| HNRNPC | HepG2 | ENCFF440ROZ |
| HNRNPC | K562 | ENCFF167CDB |
| HNRNPK | HepG2 | ENCFF855CPQ |
| HNRNPK | K562 | ENCFF918XJQ |
| HNRNPL | HepG2 | ENCFF266TKW |
| HNRNPL | K562 | ENCFF917CBK |
| HNRNPM | HepG2 | ENCFF752JNY |
| HNRNPM | K562 | ENCFF445ENC |
| HNRNPU | HepG2 | ENCFF039XQD |
| HNRNPU | K562 | ENCFF241DVU |

See next page

| Protein | Cell line | eCLIP ENCODE accession |
|---|---|---|
| HNRNPUL1 | HepG2 | ENCFF610JVZ |
| HNRNPUL1 | K562 | ENCFF889AWX |
| IGF2BP1 | HepG2 | ENCFF442USD |
| IGF2BP1 | K562 | ENCFF650LMV |
| IGF2BP2 | K562 | ENCFF524ZZB |
| IGF2BP3 | HepG2 | ENCFF886SDQ |
| ILF3 | HepG2 | ENCFF071QDP |
| ILF3 | K562 | ENCFF385MJU |
| KHDRBS1 | K562 | ENCFF027UTH |
| KHSRP | HepG2 | ENCFF771QAU |
| KHSRP | K562 | ENCFF031FMO |
| LARP4 | HepG2 | ENCFF534JCV |
| LARP4 | K562 | ENCFF947CIN |
| LARP7 | HepG2 | ENCFF307MDW |
| LARP7 | K562 | ENCFF809CKN |
| LIN28B | HepG2 | ENCFF341XMP |
| LIN28B | K562 | ENCFF061XNA |
| LSM11 | HepG2 | ENCFF985ECA |
| LSM11 | K562 | ENCFF858EHN |
| MATR3 | HepG2 | ENCFF587KKM |
| MATR3 | K562 | ENCFF246EPM |
| METAP2 | K562 | ENCFF644LPB |
| MTPAP | K562 | ENCFF883KCN |
| NCBP2 | HepG2 | ENCFF692RZM |
| NCBP2 | K562 | ENCFF886MLH |
| NIP7 | HepG2 | ENCFF033MGP |
| NIPBL | K562 | ENCFF554NIV |
| NKRF | HepG2 | ENCFF045HSV |
| NOL12 | HepG2 | ENCFF914WFQ |

See next page

**Supplementary Table S1** – continued from previous page

| Protein | Cell line | eCLIP ENCODE accession |
|---|---|---|
| NOLC1 | HepG2 | ENCFF137YFD |
| NOLC1 | K562 | ENCFF327YTD |
| NONO | K562 | ENCFF730QRI |
| NPM1 | K562 | ENCFF154AYH |
| NSUN2 | K562 | ENCFF233PRB |
| PABPC4 | K562 | ENCFF452MJL |
| PABPN1 | HepG2 | ENCFF709FEU |
| PCBP1 | HepG2 | ENCFF098REE |
| PCBP1 | K562 | ENCFF900IOH |
| PCBP2 | HepG2 | ENCFF642GNE |
| PHF6 | K562 | ENCFF588ITW |
| POLR2G | HepG2 | ENCFF591BUK |
| PPIG | HepG2 | ENCFF986XOC |
| PPIL4 | K562 | ENCFF559HGK |
| PRPF4 | HepG2 | ENCFF227EJF |
| PRPF8 | HepG2 | ENCFF048YPA |
| PRPF8 | K562 | ENCFF858UKE |
| PTBP1 | HepG2 | ENCFF726SQU |
| PTBP1 | K562 | ENCFF907HNN |
| PUM1 | K562 | ENCFF094MQV |
| PUM2 | K562 | ENCFF880MWQ |
| PUS1 | K562 | ENCFF247NXL |
| QKI | HepG2 | ENCFF704OCI |
| QKI | K562 | ENCFF786UOW |
| RBFOX2 | HepG2 | ENCFF871NYM |
| RBFOX2 | K562 | ENCFF206RIM |
| RBM15 | HepG2 | ENCFF054VLU |
| RBM15 | K562 | ENCFF597MMG |
| RBM22 | HepG2 | ENCFF293IZG |

See next page

**Supplementary Table S1** – continued from previous page

| Protein | Cell line | eCLIP ENCODE accession |
|---|---|---|
| RBM22 | K562 | ENCFF972ZMJ |
| RBM5 | HepG2 | ENCFF927KRA |
| RPS11 | K562 | ENCFF313WDF |
| RPS3 | HepG2 | ENCFF301IJW |
| RPS3 | K562 | ENCFF530HTL |
| SAFB | HepG2 | ENCFF232WUE |
| SAFB | K562 | ENCFF953WTP |
| SAFB2 | K562 | ENCFF594EDV |
| SBDS | K562 | ENCFF051EEW |
| SDAD1 | HepG2 | ENCFF825UVD |
| SDAD1 | K562 | ENCFF114EUH |
| SERBP1 | K562 | ENCFF295HRZ |
| SF3A3 | HepG2 | ENCFF950VZO |
| SF3B1 | K562 | ENCFF887ARJ |
| SF3B4 | HepG2 | ENCFF073IJF |
| SF3B4 | K562 | ENCFF649GEE |
| SFPQ | HepG2 | ENCFF139NXB |
| SLBP | K562 | ENCFF623WGE |
| SLTM | HepG2 | ENCFF121RVH |
| SLTM | K562 | ENCFF696QWE |
| SMNDC1 | HepG2 | ENCFF736RYV |
| SMNDC1 | K562 | ENCFF943XOU |
| SND1 | HepG2 | ENCFF609LWQ |
| SND1 | K562 | ENCFF211TRO |
| SRSF1 | HepG2 | ENCFF934ANS |
| SRSF1 | K562 | ENCFF886XUO |
| SRSF7 | HepG2 | ENCFF317HGW |
| SRSF7 | K562 | ENCFF780OCY |
| SRSF9 | HepG2 | ENCFF765PIF |

See next page

| Protein | Cell line | eCLIP ENCODE accession |
|---|---|---|
| SSB | HepG2 | ENCFF848JWA |
| SSB | K562 | ENCFF328KJW |
| STAU2 | HepG2 | ENCFF678FCX |
| SUB1 | HepG2 | ENCFF335JTC |
| SUGP2 | HepG2 | ENCFF191XRG |
| SUPV3L1 | HepG2 | ENCFF999ZAW |
| SUPV3L1 | K562 | ENCFF239GPP |
| TAF15 | HepG2 | ENCFF566EAE |
| TAF15 | K562 | ENCFF822NWY |
| TARDBP | K562 | ENCFF037TVC |
| TBRG4 | HepG2 | ENCFF869RUC |
| TBRG4 | K562 | ENCFF618WYU |
| TIA1 | HepG2 | ENCFF759KCD |
| TIA1 | K562 | ENCFF918KMT |
| TIAL1 | HepG2 | ENCFF612HOP |
| TRA2A | HepG2 | ENCFF766OCH |
| TRA2A | K562 | ENCFF726PFJ |
| TROVE2 | HepG2 | ENCFF611SOO |
| TROVE2 | K562 | ENCFF794RJQ |
| U2AF1 | HepG2 | ENCFF056LEW |
| U2AF1 | K562 | ENCFF640IHY |
| U2AF2 | HepG2 | ENCFF721PWF |
| U2AF2 | K562 | ENCFF290DFO |
| UCHL5 | HepG2 | ENCFF573DXD |
| UCHL5 | K562 | ENCFF365LFU |
| UPF1 | HepG2 | ENCFF687EQE |
| UPF1 | K562 | ENCFF597NVR |
| UTP18 | HepG2 | ENCFF644EIR |
| UTP18 | K562 | ENCFF783INO |

See next page

**Supplementary Table S1** – continued from previous page

| Protein | Cell line | eCLIP ENCODE accession |
|---|---|---|
| UTP3 | K562 | ENCFF028GAJ |
| WDR3 | K562 | ENCFF560RWD |
| WDR43 | HepG2 | ENCFF945NFS |
| WDR43 | K562 | ENCFF244JER |
| WRN | K562 | ENCFF404HHM |
| XPO5 | HepG2 | ENCFF327VCE |
| XRCC6 | HepG2 | ENCFF993VRZ |
| XRCC6 | K562 | ENCFF958GMP |
| XRN2 | HepG2 | ENCFF321HPH |
| XRN2 | K562 | ENCFF695OLO |
| YBX3 | HepG2 | ENCFF185OEI |
| YBX3 | K562 | ENCFF746XLE |
| YWHAG | K562 | ENCFF932UXC |
| ZC3H11A | HepG2 | ENCFF592WIH |
| ZC3H11A | K562 | ENCFF258HOX |
| ZC3H8 | K562 | ENCFF382URH |
| ZNF622 | K562 | ENCFF285IGP |
| ZNF800 | HepG2 | ENCFF343TCO |
| ZNF800 | K562 | ENCFF105HMJ |
| ZRANB2 | K562 | ENCFF058OOW |

**Supplementary Table S2.** Extended RNA-binding domains list used for this study.

| Domain mnemonic |
|---|
| NHL |
| SAM_1 |
| KH_1 |
| KH_2 |
| CSD |
| DSRM |

See next page

| **Domain mnemonic** |
|---|
| La |
| PUF |
| S1 |
| YTH |
| zf-CCCH |
| zf-CCHC |
| zf-CCHH |
| zf-RanBP |
| DEAD |
| Helicase_C |
| Piwi |
| Tudor |
| OB |
| G-patch |
| Pumilio_1 |
| RNase_H |
| RNase_P |
| TruB_N |
| LARP_1 |
| U2AF |
| SF1 |
| PTB |
| hnRNP |

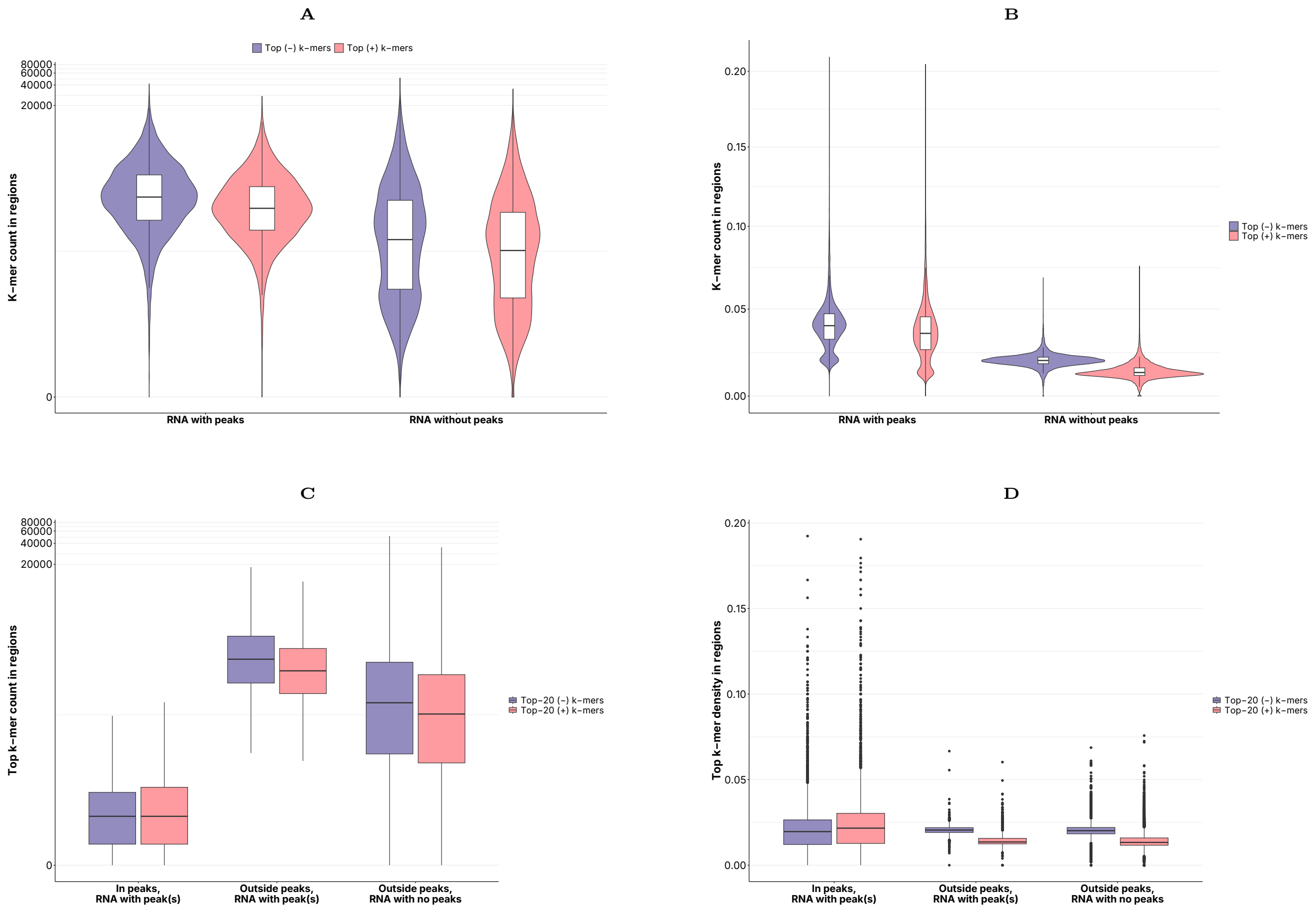


**Supplementary Figure S1.** Stratified analysis of DDX42 model k-mer importance correlation with the k-mer being the statistically significant peak of the DDX42-RNA interaction: **A** figure from original manuscript; **B** same, but the number of important k-mers was normalized to total region length on a transcript; **C** number of important k-mers on transcripts with or without peaks on them, if and RNA has a peak on it, it is split into peaks and regions outside the peaks; **D** same, but normalized to the according region's legth transcript-wise.

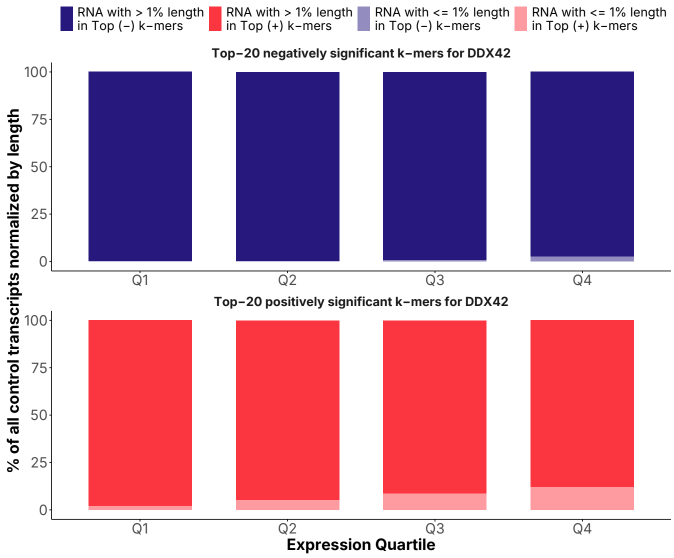


**Supplementary Figure S2.** Enrichment of significant k-mers identified by the model based for DDX42 protein in highly and lowly expressed transcripts, normalized by gene length.

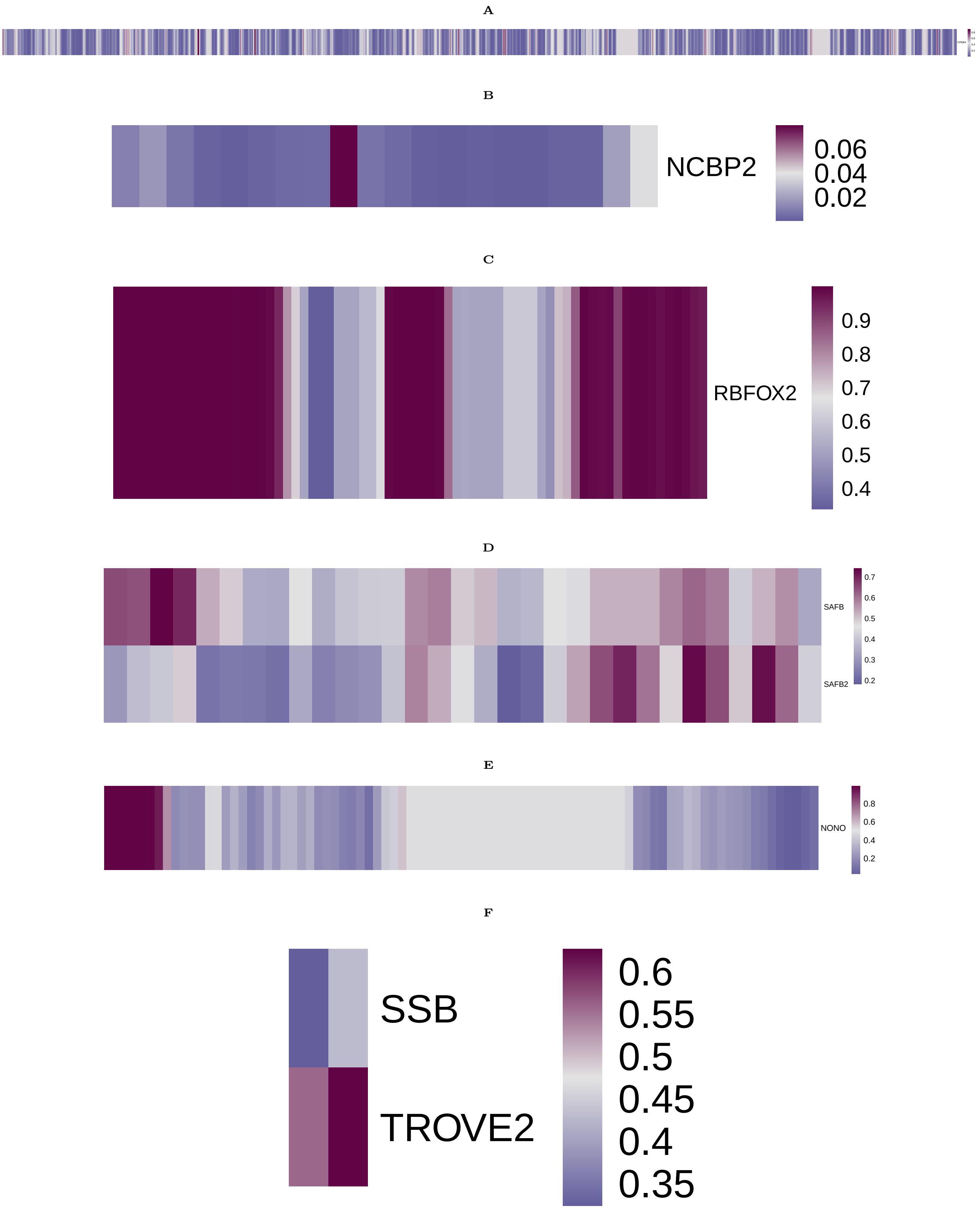


**Supplementary Figure S3.** Examples of single-protein models ncRNA-protein contact inference in PLERIO framework: **A** CPEB-DT and CPEB4 protein, possible family-wise regulation; **B** THOC7-AS1 and NCBP2 protein, possibly connected via RNA export; **C** GLIS2-AS1 and RBFOX2 protein, possibly connected through involvement in splicing control; **D** NDUFB2-AS1 RNA and SAFB and SAFB2 proteins, possible antisense RNA interaction; **E** PRKAR2A-AS1 and NONO protein, possibly interacting in paraspeckles; **F** RN7SL23P RNA and SSB and TROVE2 proteins.

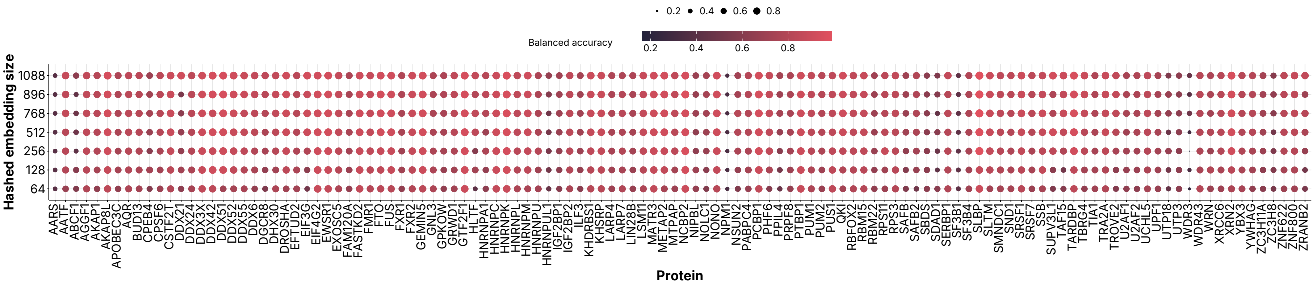


**Supplementary Figure S4.** The effects of varying k-mer hasing output vector dimension on model performance.

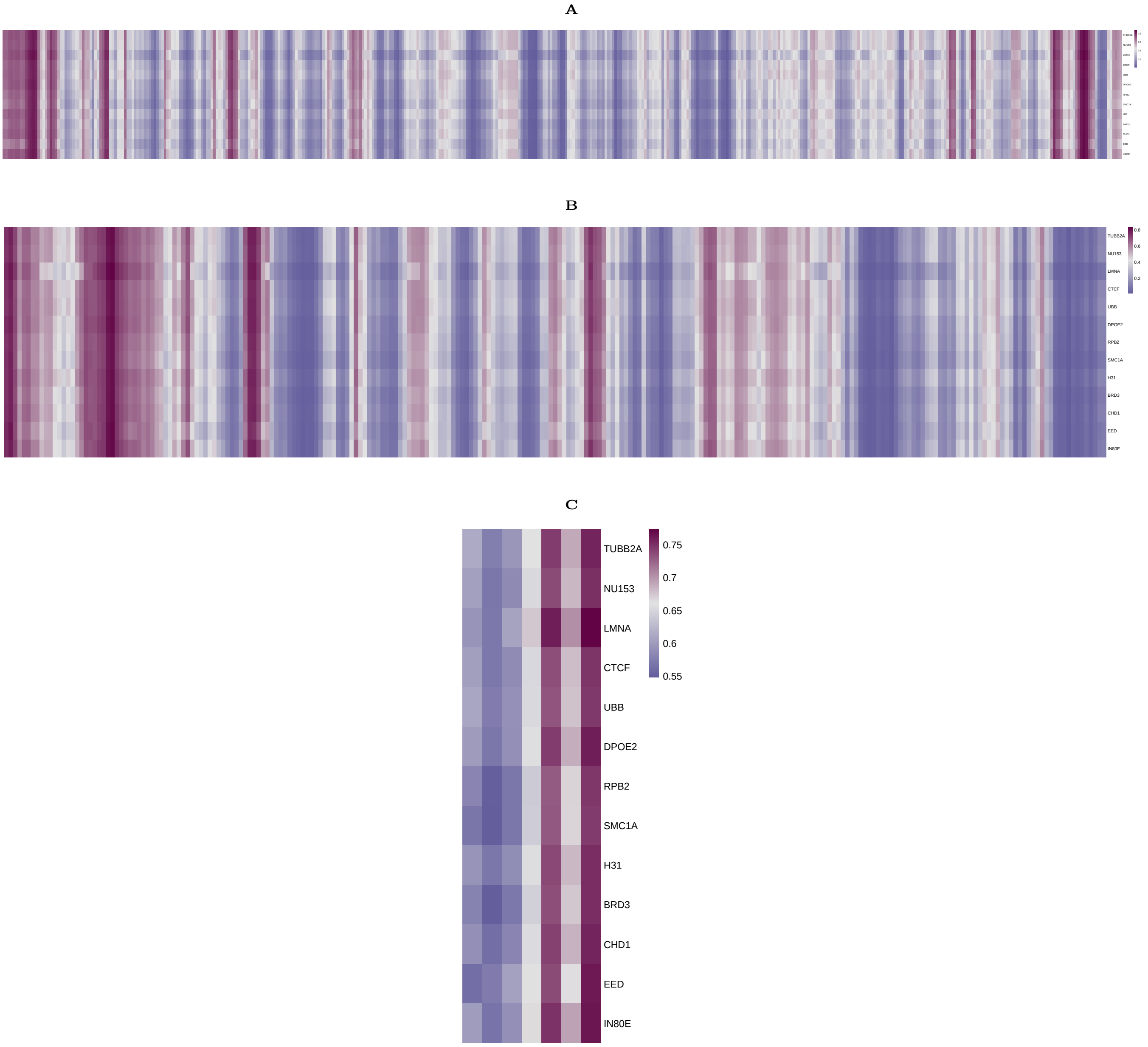


**Supplementary Figure S5.** Examples of multi-protein models ncRNA-protein contact inference in PLERIO framework: **A** NEAT1; **B** HOTAIR; **C** TERC; with multiple proteins covering a wide range of functions where RNA-binding might not be needed. Yet, the interaction profile is practically the same.

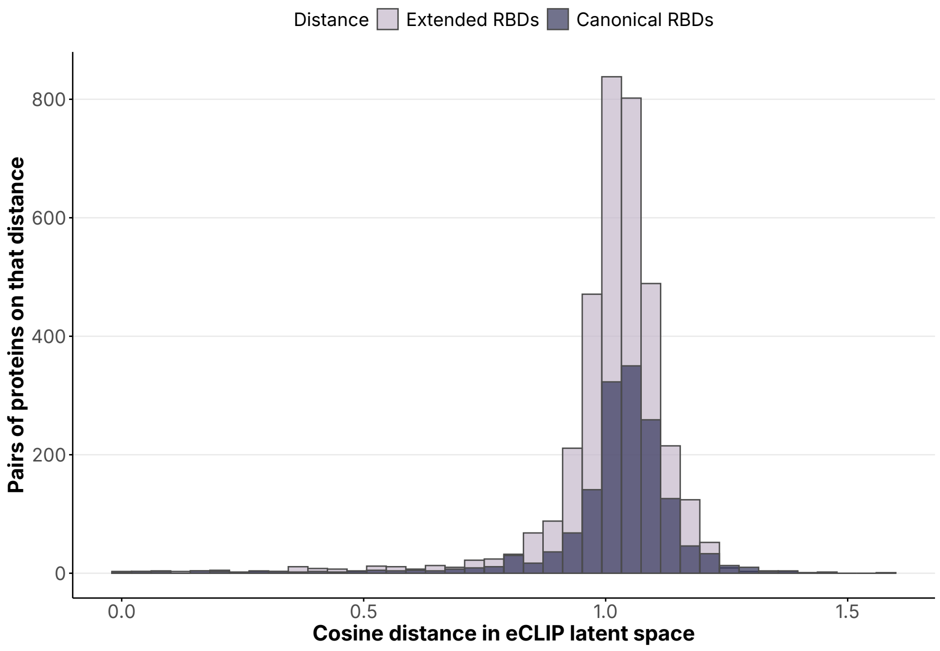


**Supplementary Figure S6.** Distribution of cosine distances in eCLIP-based latent space obtained by JPLE algorithm on available proteins with a canonical RBD or a domain that can be less strictly considered RNA-binding. Wasserstein's W1 distance between two distributions is 0.01697.
